# Proteomics reveals unique identities of human TGF-β-induced and thymus-derived CD4^+^ regulatory T cells

**DOI:** 10.1101/2021.10.30.466569

**Authors:** Mark Mensink, Ellen Schrama, Eloy Cuadrado, Maartje van den Biggelaar, Derk Amsen, Sander de Kivit, Jannie Borst

## Abstract

The CD4^+^ regulatory T (Treg) cell lineage, as defined by FOXP3 expression, comprises thymus-derived (t)Treg cells and peripherally induced (p)Treg cells. In human, naïve tTreg cells can be isolated from blood, while effector tTreg and pTreg cells cannot be purified reliably for lack of cell surface markers. As a model for Treg cells, studies often employ TGF-β-induced (i)Treg cells generated from CD4^+^ conventional T (Tconv) cells *in vitro*. Here, we describe the relationship of iTreg cells to tTreg and Tconv cells, as optimally purified from human blood. Proteomic analysis revealed that each of these cell populations has a unique protein expression pattern before and after CD3/CD28-mediated activation. iTreg cells had limited overlap in protein expression with tTreg cells and exhibited markedly differential expression of proteins involved in signal transduction and metabolism. Whereas iTreg and Tconv cells responded strongly to CD3/CD28-mediated activation, tTreg cells showed a modest response, which reflects their adaptations in signal transduction pathways. As a benchmark, we used a previously defined proteomic signature that discerns *ex vivo* naïve and effector Treg cells from Tconv cells and includes conserved Treg cell properties. This Treg cell core signature was largely absent in iTreg cells, as exemplified by STAT4 expression that may contribute to pro-inflammatory functions. In addition, we used a proteomic signature that distinguishes *ex vivo* effector Treg cells from Tconv cells and naïve Treg cells. This effector Treg cell signature was partially present in iTreg cells, indicating that they do have protein expression features in common with *ex vivo* effector Treg cells. In conclusion, iTreg cells are distinct from tTreg cells and largely lack the Treg cell core proteomic signature, indicating that caution is warranted when using iTreg cells as a model for Treg cell populations found *in vivo*.

## 1 Introduction

Immune responses depend on a balance between the opposing activities of conventional (Tconv) and regulatory T (Treg) cells (1). While Tconv cells are key mediators of adaptive immunity, Treg cells are specialized in maintaining immune tolerance and controlling inflammation (1). Based on their origin, Treg cells can be classified into two main groups. Thymus-derived (t)Treg cells are primarily directed at self-antigens and stably express the Treg cell master transcription factor FOXP3, which is indispensable for their identity and suppressive function (2–6). In addition, they express the transcription factor Helios (IKZF2), which is consequently used as a marker of stable Treg cells (7, 8). Peripherally induced (p)Treg cells are converted from Tconv cells in response to foreign antigens (9) and reportedly have less stable FOXP3 expression (10–13), in particular recently after their development (14). Stable expression of FOXP3 is established epigenetically by demethylation of several regions within the *FOXP3* gene locus, including the so-called Treg-specific demethylated region (TSDR) in the CNS2 enhancer (5, 15).

Based on expression of CD45RA, human Treg cells from peripheral blood can be divided into naïve (CD45RA^+^) and effector (CD45RA^−^) Treg cells (16). Since conversion of Tconv cells into pTreg cells requires their activation, it stands to reason that the naïve Treg cell population is mainly composed of tTreg cells (17, 18). Such naïve Treg cells can differentiate into effector Treg cells in response to inflammatory signals (19). Therefore, the effector Treg cell population arguably consists of a mixture of pTreg cells and activated tTreg cells. Amongst human effector Treg cells, it is currently not possible to definitively discriminate tTreg and pTreg cells for a lack of cell surface markers (20).

Naïve Treg cells occur in low abundance in the blood (21, 22) and effector Treg cells from the blood are not amenable to expansion *in vitro* (16, 23). For these reasons, *in vitro* Treg cell studies often employ TGF-β-induced (i)Treg cells that are generated from Tconv cells (24–27). Such iTreg cells resemble Treg cells by their expression of FOXP3 and their suppressive capacity (24, 25). However, in terms of FOXP3 expression and suppressive function, iTreg cells are less stable than tTreg cells. For example, adoptively transferred iTreg cells in mice lost Foxp3 expression and failed to prevent graft-versus-host disease, in contrast to tTreg cells (28). Similarly, others showed that iTreg cells had reduced Foxp3 expression and suppressive capacity after removal of TGF-β *in vitro* and that the majority of iTreg cells rapidly lost Foxp3 expression following adoptive transfer in wild-type mice (29).

It is unclear whether iTreg cells represent an equivalent of *ex vivo* Treg cells. We recently described that *ex vivo* naïve and effector Treg cells from human peripheral blood have a common proteomic Treg cell signature, which sets them apart from other CD4^+^ T cell subsets (22). This Treg cell core signature reflects unique Treg cell properties regarding signal transduction, transcriptional regulation and cell metabolism. In addition, we identified a proteomic signature that distinguishes *ex vivo* effector Treg cells from both Tconv cells and naïve Treg cells. Using these proteomic signatures as benchmarks, we compared in the current study the proteomes of iTreg, tTreg and Tconv cells to determine to what extent *in vitro*-generated iTreg cells resemble *ex vivo* Treg cells.

## 2 Methods

### 2.1 Cell isolation and flow cytometric sorting

Human materials were obtained in compliance with the Declaration of Helsinki and the Dutch rules regarding the use of human materials from voluntary donors. Buffy coats from healthy anonymized male donors were obtained after their written informed consent, as approved by the internal ethical board of Sanquin, the Netherlands. Total CD4^+^ T cells were collected from buffy coats by first performing a Ficoll-Paque Plus (GE Healthcare) density gradient centrifugation to isolate peripheral blood mononuclear cells (PBMC) and subsequently using CD4 magnetic MicroBeads (Miltenyi Biotec) according to the manufacturers’ protocols. Alternatively, total CD4^+^ T cells were directly isolated from buffy coats using the StraightFrom Buffy Coat CD4 MicroBead kit (Miltenyi Biotec) according to the manufacturer’s instructions. To sort naïve Tconv and tTreg cells, CD4^+^ T cells were stained with CD4-PE-Cy7, CD127-BV421 (BioLegend), CD25-PE (BD Biosciences), CD45RA-FITC (Immunotools) and GPA33-AF647 (30) monoclonal antibodies (mAbs) as indicated (**Supplementary Figure 1**). Cells were sorted on a MoFlo Astrios using Summit software version 6.2 (Beckman Coulter) or BD FACS Aria II using FACSDiva software version 8 (BD Biosciences). Either propidium iodide (Sigma) or near-IR dead cell stain kit (Invitrogen) was used as a live/dead marker.

### 2.1 T-cell expansion cultures and restimulation

Following cell sorting, human naïve Tconv and tTreg cells were cultured in 96-well round bottom plates (Greiner) at 1 × 10^4^ cells per well using IMDM (Gibco, Life Technologies) supplemented with 8% FCS (Sigma), penicillin/streptomycin (Roche) and 300 IU/ml IL-2 (DuPont Medical), hereinafter referred to as T-cell medium, at 37°C and 5% CO_2_. Cells were expanded using agonistic mAbs against CD3 (clone CLB-T3/4.E, IgE isotype, Sanquin, 0.1 μg/ml) and CD28 (clone CLB-CD28/1, Sanquin, 0.2 μg/ml) in solution. After four days, fresh T-cell medium was added. From day 7 to day 14, cells were cultured in 24-well plates (Greiner) at 5 × 10^5^ cells per ml, while fresh T-cell medium was added on day 11. On day 14, cells were cultured in 6-well plates (Corning) at 1 × 10^6^ cells per ml for four days using fresh T-cell medium in the absence of agonistic mAbs. To generate iTreg cells, naïve Tconv cells were cultured as described above in the presence of TGF-β (Peprotech, 10 ng/ml). Prior to restimulation experiments, dead cells were removed by Ficoll-Paque Plus density gradient centrifugation. T cells were restimulated as indicated using T-cell medium in the presence of agonistic mAbs against CD3 (0.1 μg/ml) and CD28 (0.2 μg/ml), but without TGF-β.

### 2.2 Flow cytometry

For expression analysis of cell surface molecules, T cells were washed in PBS/1% FCS and stained using the following mAbs in appropriate combinations: anti-OX40-PE-Cy7 (BioLegend), anti-GITR- BV421 (BioLegend), anti-PD-1-APC-Cy7 (BioLegend), anti-CD39-BV510 (BioLegend), anti-ITGA4- PE (CD49d) (BD Biosciences), anti-ICOS-PerCP-Cy5.5 (BioLegend), anti-CCR4-PE-Cy7 (BioLegend), anti-HLA-DR-APC-eFluor 780 (Invitrogen), anti-HLA-DR-BV605 (BD Biosciences) and anti-FAS-BB700 (CD95) (BD Biosciences). Cell surface expression of CTLA-4 was measured by adding anti-CTLA-4-PE-Dazzle594 mAb (BioLegend) in culture during restimulation for 24 h. Near-IR dead cell stain kit (Invitrogen) was used as a live/dead marker. For intracellular staining, cells were fixed and permeabilized using the FOXP3 transcription factor staining buffer set (Invitrogen) according to the manufacturer’s instructions. Next, cells were stained with combinations of anti-FOXP3-APC (Invitrogen), anti-Helios-PE-Cy7, anti-CTLA-4-PE-Dazzle594 (BioLegend) and anti-NFATC2-AF488 (Cell Signaling Technology) mAbs. Flow cytometry was performed using a BD LSR Fortessa or BD LSR II cell analyzer (BD Biosciences). Sample acquisition was performed using FACSDiva software version 8 and data were analyzed using FlowJo software version 10.6.0.

### 2.3 Methylation assay of the TSDR

The methylation status of the TSDR within the *FOXP3* locus was determined in Tconv, tTreg and iTreg cells as described previously (31). Only male donors were used for these analyses, given the X-chromosomal location of the *FOXP3* locus. In brief, expanded T cells were resuspended in PBS for proteinase K digestion, followed by bisulfite conversion of DNA using the EZ DNA Methylation-Direct kit (Zymo Research) and methylation-specific quantitative PCR using iQTM SYBR Green Supermix (Bio-Rad) (32). Methylation-specific primers were 5’-CGATAGGGTAGTTAGTTTTCGGAAC-3’ and 5’- CATTAACGTCATAACGACCGAA-3’. Demethylation-specific primers were 5’-TAGGGTAGT- TAGTTTTTGGAATGA-3’ and 5’-CCATTAACATCATAACAACCAAA-3’. Melt curve analysis was performed on a LightCycler® 480-II (Roche). Methylation of the TSDR (%) was calculated using the following formula: 100/(1+2^Ct[CG]-Ct[TG]^), where Ct[CG] is defined as Ct values obtained using methylation-specific primers and Ct[TG] is defined as Ct values obtained using demethylation-specific primers.

### 2.4 Suppression assay

Total human PBMCs were washed and resuspended in PBS, followed by incubation for 8 min using 5 μM CellTrace Violet dye (Invitrogen). Following fluorescent labeling, an equal volume of FCS was added and cells were subsequently washed twice in IMDM supplemented with 8% FCS. CellTrace Violet-labeled PBMCs were cocultured with expanded Tconv, tTreg or iTreg cells for 96 h at indicated ratios, in the presence of anti-CD3 mAb (0.05 μg/ml). Proliferation of PBMCs was examined by tracing CellTrace Violet dye dilution by flow cytometry on a BD LSR Fortessa or BD LSR II cell analyzer. Data were analyzed using FlowJo software (version 10.6.0).

### 2.5 Proteomics

Proteomics data were generated as described before by Cuadrado et al. (22), including analysis of expanded Tconv, tTreg and iTreg cells. In short, at least 1 × 10^6^ expanded T cells were restimulated for 24 h using T-cell medium in the presence of agonistic mAbs against CD3 (0.1 μg/ml) and CD28 (0.2 μg/ml) or not. Unstimulated and restimulated cells were washed, lysed and processed for mass spectrometry. RAW mass spectrometry files were processed previously according to Cuadrado et al. (22), here followed by the removal of potential contaminants and reverse hits using Perseus (version 1.6.12). Label-free quantification (LFQ) values were log_2_-transformed and the biological replicates were grouped based on cell type. Proteins with a minimum of 5 valid values in at least one cell type were included for further analysis. For analysis of the effector Treg cell signature, a minimum of 4 valid values in at least one cell type was used. Missing value imputation was performed by replacing missing values by random numbers drawn from the lower part of the normal distribution (width = 0.3, shift = 1.8). Next, proteomics data were analyzed by principal component analysis (PCA), ANOVA and Student’s *t*-test using Qlucore Omics Explorer (version 3.8). After z-score normalization, clusters of proteins with similar expression patterns were identified using unsupervised hierarchical clustering and visualized in heat maps. Protein clusters were subjected to overrepresentation analysis to assess enrichment of Gene Ontology (GO) biological processes using the STRING app (version 1.6.0) in Cytoscape (version 3.9.0). To refine the analysis of biological processes, a redundancy cutoff was applied when significant GO terms contained large overlap based on enriched genes. The proteomic response to restimulation was analyzed for each cell type using GO biological processes in gene set enrichment analysis (GSEA) software (version 4.1.0). GSEA was performed using default parameters and gene set permutations. To identify related groups of biological processes that were differentially regulated following restimulation of T cells, the EnrichmentMap app (version 3.3.2) was used in Cytoscape. The largest gene set clusters were displayed using manually assigned annotations that describe the general biological processes represented by the clusters. To assess commonalities and differences between the proteomic responses of the studied cell populations, the DiVenn tool (33) was used.

### 2.6 Statistical analysis

Data were analyzed using GraphPad Prism (version 9.0.1), except for proteomics data. Statistical analyses were performed as indicated in the figure legends. Log_2_ transformation was performed when data were not normally distributed. Data are presented as mean ± SEM and a two-sided *p* < 0.05 was considered statistically significant.

## 3 Results

### 3.1 Validation of Tconv, tTreg and iTreg cell populations used in this study

To obtain the cell populations of interest, we first isolated CD4^+^ T cells from human peripheral blood. These cells were flow cytometrically sorted based on a CD25^lo^CD127^hi^CD45RA^+^GPA33^int^ phenotype to obtain naïve Tconv cells, or a CD25^hi^CD127^lo^CD45RA^+^GPA33^hi^ phenotype to obtain naïve Treg cells, as described before (21, 31) (**Supplementary Figure 1**). After sorting, the cells were expanded using agonistic mAbs against CD3 and CD28 in the presence of IL-2. Cells derived from naïve Treg cells are hereinafter referred to as tTreg cells. To generate iTreg cells, TGF-β was added to Tconv cell cultures. After two weeks of expansion *in vitro*, cells were “rested” for four days prior to restimulation with anti-CD3/CD28 antibodies for 24 h and analysis (**Figure 1A**).

**Figure 1.**
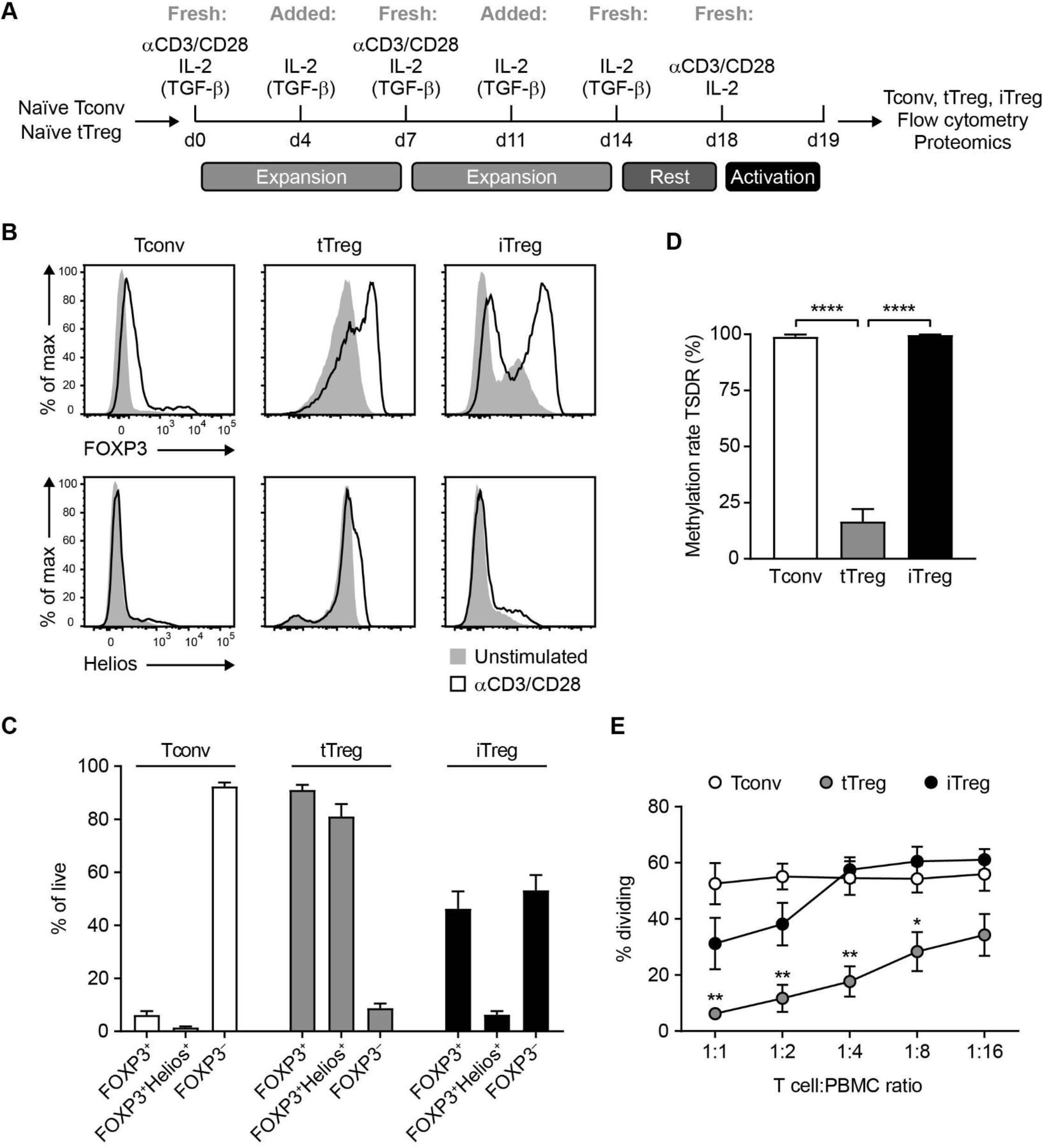
Characterization of Tconv, tTreg and iTreg cell populations. **(A)** Schematic overview of expansion protocols for Tconv, tTreg and iTreg cells. TGF-β was used to generate iTreg cells. Prior to analysis, expanded cells were restimulated for 24 h or not, as indicated in figure legends. **(B)** Flow cytometric analysis of FOXP3 and Helios (IKZF2) protein levels in Tconv, tTreg and iTreg cells at 24 h after restimulation with anti-CD3/CD28 mAbs or unstimulated control (representative of *n* = 7). **(C)** Quantification of the frequency of FOXP3^+^, FOXP3^+^Helios^+^ and FOXP3^−^ cells in the resting Tconv, tTreg and iTreg cell populations after expansion (*n* = 7). **(D)** Analysis of the methylation status of the TSDR in the *FOXP3* locus in Tconv, tTreg and iTreg cells. One-way ANOVA with Tukey’s *post hoc* test was used for statistical analysis (*n* = 6-7). **(E)** Assessment of the suppressive capacity of Tconv, tTreg and iTreg cells that were cocultured for four days with CellTrace Violet-labeled responder PBMCs in the presence of agonistic mAb to CD3. The percentage of dividing PBMCs is displayed (*n* = 4). One-way ANOVA with Bonferroni’s *post hoc* test was used for statistical analysis (*n* = 4). **(C-E)** Data are presented as mean ± SEM. Sample size (*n*) represents cells from individual donors, analyzed in independent experiments. \**p* < 0.05, \*\**p* < 0.01, \*\*\*\**p* < 0.0001.

At this point, Tconv cells were FOXP3^−^Helios^−^ and largely retained this phenotype after CD3/CD28-mediated restimulation (**Figure 1B, C**). tTreg cells showed a uniform expression of FOXP3 and Helios (IKFZ2), both before and after restimulation, suggesting that the isolation and cell culture method generated pure tTreg cells as based on Helios expression (**Figure 1B, C**). The iTreg cells showed bimodal FOXP3 expression, with an upregulation of FOXP3 upon CD3/CD28-mediated restimulation, while Helios was minimally expressed, as described (7) (**Figure 1B, C**).

The TSDR of the *FOXP3* gene locus had the highly demethylated configuration in tTreg cells and was methylated in Tconv and iTreg cells, in agreement with published data (5, 34) (**Figure 1D**). Next, we functionally characterized the cell populations in a standard assay, testing their ability to suppress CD3-induced proliferation of CellTrace Violet-labeled Tconv cells from peripheral blood. In this assay, tTreg cells were suppressive and more so than iTreg cells, while Tconv cells were not, in agreement with earlier research (35) (**Figure 1E**). Together, these data validate the purification and cell culture protocols to generate tTreg, iTreg and Tconv cell populations for further characterization.

### 3.2 Expression of characteristic cell surface receptors on tTreg and iTreg cells

We next examined the expression of hallmark cell surface receptors that are associated with Treg cells. Among these, CTLA-4 is a key mediator of suppression by tTreg cells (36) by reducing the availability of CD80/86 for the costimulatory receptor CD28 on Tconv cells (37, 38). As CTLA-4 recycles between the cell surface and endosomes (39), we measured both total (cell surface plus intracellular) CTLA-4 levels, as well as specifically the CTLA-4 levels on the cell surface only. Both tTreg and iTreg cells showed higher total CTLA-4 expression than Tconv cells (**Figure 2A**, **Supplementary Figure 2**). All three cell types upregulated total CTLA-4 expression upon CD3/CD28- mediated restimulation, but levels were still higher in tTreg and iTreg cells than in Tconv cells. CTLA- 4 was only found on the cell surface of expanded cells after CD3/CD28-mediated restimulation, with tTreg cells showing the highest levels (**Figure 2A**, **Supplementary Figure 2**). The costimulatory receptors OX40 and GITR have received attention as Treg cell markers and mediators of Treg cell function (40–42). These two receptors were expressed by all expanded cell types, but only following CD3/CD28-mediated restimulation, with highest levels on tTreg cells (**Figure 2A**, **Supplementary Figure 2**). Upon CD3/CD28-mediated restimulation, both iTreg and Tconv cells but not tTreg cells upregulated PD-1, which is a co-inhibitory receptor for both Tconv and Treg cells (43) (**Figure 2A**, **Supplementary Figure 2**). We also analyzed CD39 (ENTPD1), which facilitates extracellular ATP hydrolysis that generates adenosine, a molecule with immunosuppressive effects on Tconv cells (44). CD39 protein levels were highest on iTreg cells both before and after restimulation (**Figure 2A**, **Supplementary Figure 2**). These data show that tTreg and iTreg cells exhibit different expression patterns of these Treg cell-associated cell surface molecules (**Figure 2B**).

**Figure 2.**
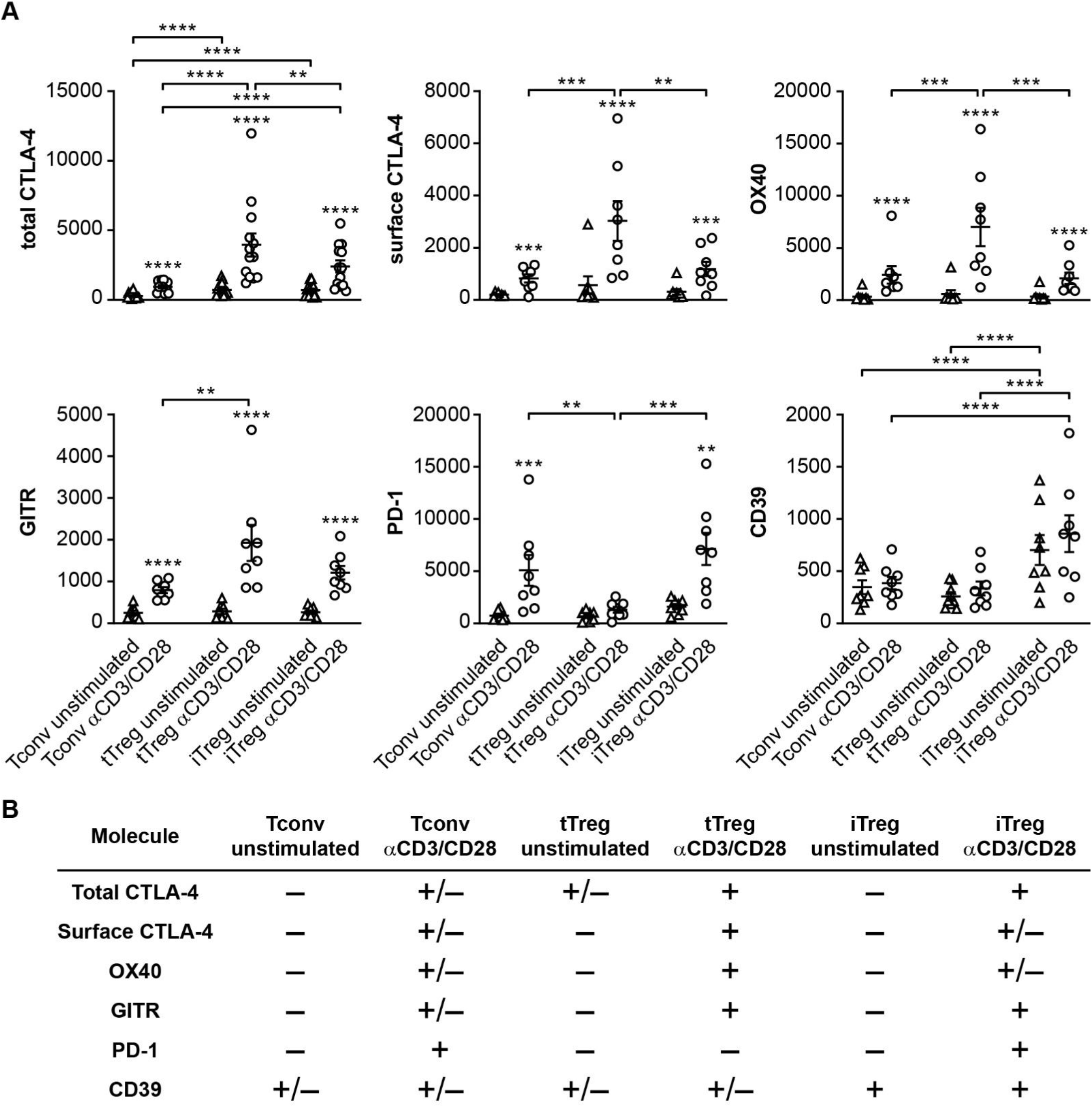
Expression of characteristic cell surface molecules associated with Treg cells. **(A)** Quantification of flow cytometric analysis of the protein expression of total CTLA-4 (cell surface plus intracellular) (*n* = 13), surface CTLA-4 (*n* = 8), surface OX40 (*n* = 8), surface GITR (*n* = 8), surface PD-1 (*n* = 8) and surface CD39 (ENTPD1) (*n* = 8) on indicated cell populations. Mean fluorescence intensity (MFI) is depicted on the y-axis. Two-way ANOVA with Tukey’s *post hoc* test was used for statistical analysis. Data are presented as mean ± SEM. Sample size (*n*) represents cells from individual donors, analyzed in independent experiments. Asterisks without lines indicate comparisons between cells that were restimulated or not. \**p* < 0.05, \*\**p* < 0.01, \*\*\**p* < 0.001, \*\*\*\**p* < 0.0001. **(B)** Table summarizing the data in **A**, indicating relatively low (–), intermediate (+/–) or high (+) expression levels.

### 3.3 Proteomic analysis reveals that iTreg cells are distinct from tTreg and Tconv cells

We performed label-free quantitative proteomics to compare overall protein expression profiles in resting Tconv, tTreg and iTreg cells in an unbiased manner. Based on the levels of the 4224 proteins that were identified (**Supplementary Table 1**), PCA led to distinct grouping of expanded Tconv, tTreg and iTreg cells (**Figure 3A**). PC1 explained most of the differences between the populations (32%). It revealed that iTreg and tTreg cells were distinct from one another and each more similar to Tconv cells. In contrast, PC2 (16%) indicated commonalities between iTreg and tTreg cells that set them apart from Tconv cells. Unsupervised hierarchical clustering, based on the 1290 proteins that are differentially expressed between the three cell types (shown as a heat map of z-scores), also revealed marked differences (**Figure 3B**, **Supplementary Table 2**). Each cell population exhibited a unique pattern of clusters of proteins with relatively high or low expression. This unsupervised analysis clustered iTreg and Tconv cells together in one branch of the dendrogram, whereas tTreg cells clustered in a different branch, showing that iTreg cells are overall more related to Tconv cells than to tTreg cells. In fact, only two small clusters of proteins were shared by iTreg and tTreg cells but not Tconv cells: cluster 1 and cluster 6, which represented high or low expression of proteins compared to Tconv cells, respectively. GO analysis indicated an overrepresentation of proteins associated with protein folding and endoplasmic reticulum stress in these two clusters (**Figure 3C**, **Supplementary Figure 3**). Notable were the shared high expression of Ikaros family member IKZF3 (Aiolos), a cofactor for FOXP3-mediated transcriptional repression (45), and the shared low expression of YY1, which inhibits FOXP3 expression and function (46).

**Figure 3.**
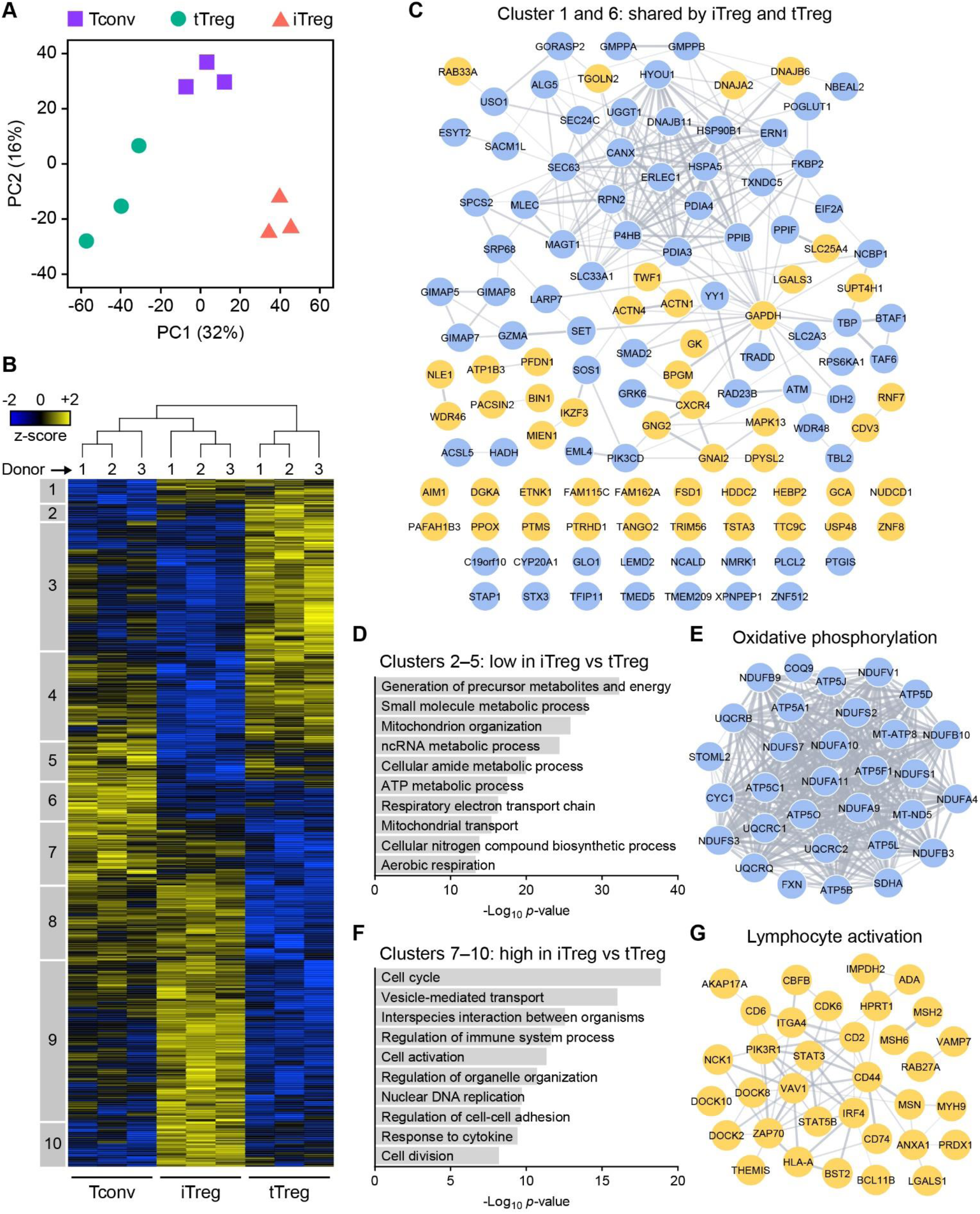
The iTreg, tTreg and Tconv cell proteomes are distinct. **(A)** PCA plot of the proteome of Tconv, tTreg and iTreg cells without restimulation (*n* = 3). **(B)** Heat map showing hierarchical clustering of the 1290 proteins that were differentially expressed between unstimulated Tconv, tTreg and iTreg cells (ANOVA, *p* < 0.05). Z-scores showing relative protein expression values are color-coded. Numbered boxes (1–10) indicate clusters of proteins. **(C)** STRING network of proteins that have shared high (cluster 1) or low (cluster 6) expression in iTreg and tTreg cells, relative to Tconv cells. Proteins with high or low expression are depicted as yellow or blue nodes, respectively. Protein associations according to STRING are connected with lines. **(D)** GO enrichment analysis of clusters 2–5 containing proteins with low expression in iTreg cells compared to tTreg cells, showing ten significantly enriched GO biological processes. **(E)** STRING network of proteins from clusters 2–5 that are involved in oxidative phosphorylation. Blue nodes indicate low expression in iTreg cells relative to tTreg cells. **(F)** GO enrichment analysis of clusters 7–10 containing proteins with high expression in iTreg cells compared to tTreg cells, showing ten significantly enriched GO biological processes. **(G)** STRING network of proteins from clusters 7–10 that are involved in lymphocyte activation. Yellow nodes indicate high expression in iTreg cells relative to tTreg cells.

Apart from this limited overlap in protein expression between iTreg and tTreg cells, a far greater number of proteins (clusters 2–5 and 7–10) was differentially expressed between these cell types, emphasizing their dissimilarity (**Figure 3B**). Proteins in clusters 2–5 were expressed at lower levels in iTreg cells than in tTreg cells and were mainly associated with metabolic processes (**Figure 3D**). Part of these clusters showed shared low protein expression by iTreg and Tconv cells (clusters 2–3), whereas the other part showed low protein expression uniquely in iTreg cells (clusters 4–5) (**Supplementary Figure 3**). Noteworthy was the lower expression of 31 proteins involved in oxidative phosphorylation in iTreg cells compared to tTreg cells (**Figure 3E**). Of these, 23 proteins were located in cluster 3, indicating that this lower expression was largely shared with Tconv cells. Clusters 7–10 consisted of proteins that were present at higher levels in iTreg cells than in tTreg cells and were involved in signal transduction, cell division, vesicle-mediated transport and cell adhesion (**Figure 3F**). These clusters included proteins related to lymphocyte activation, such as VAV1, ZAP70, STAT3 and STAT5B (**Figure 3G**). Part of these clusters indicated high protein expression shared by iTreg and Tconv cells (clusters 7–8), while the other clusters indicated high protein expression uniquely in iTreg cells (clusters 9–10) (**Supplementary Figure 3**). This analysis indicates that iTreg cells display a protein expression profile related to that of Tconv cells that largely sets them apart from tTreg cells, including metabolic features and potential responses to activation signals. However, iTreg cells do share some features with tTreg cells, in particular regarding endoplasmic reticulum-associated processes.

### 3.4 iTreg and tTreg cells respond differentially to CD3/CD28 stimulation

We also analyzed the proteome of Tconv, tTreg and iTreg cells following CD3/CD28-mediated restimulation. Distinct clusters were identified in a PCA, showing that particularly Tconv and iTreg cells altered their protein expression upon activation (**Figure 4A**). A much smaller response was observed in tTreg cells, which clustered adjacent to each other regardless of stimulation. GSEA emphasized the differential responses between the cell types (**Supplementary Figure 4A**), indicating that iTreg cells, like Tconv cells and unlike tTreg cells, strongly responded, particularly by processes associated with cell proliferation.

**Figure 4.**
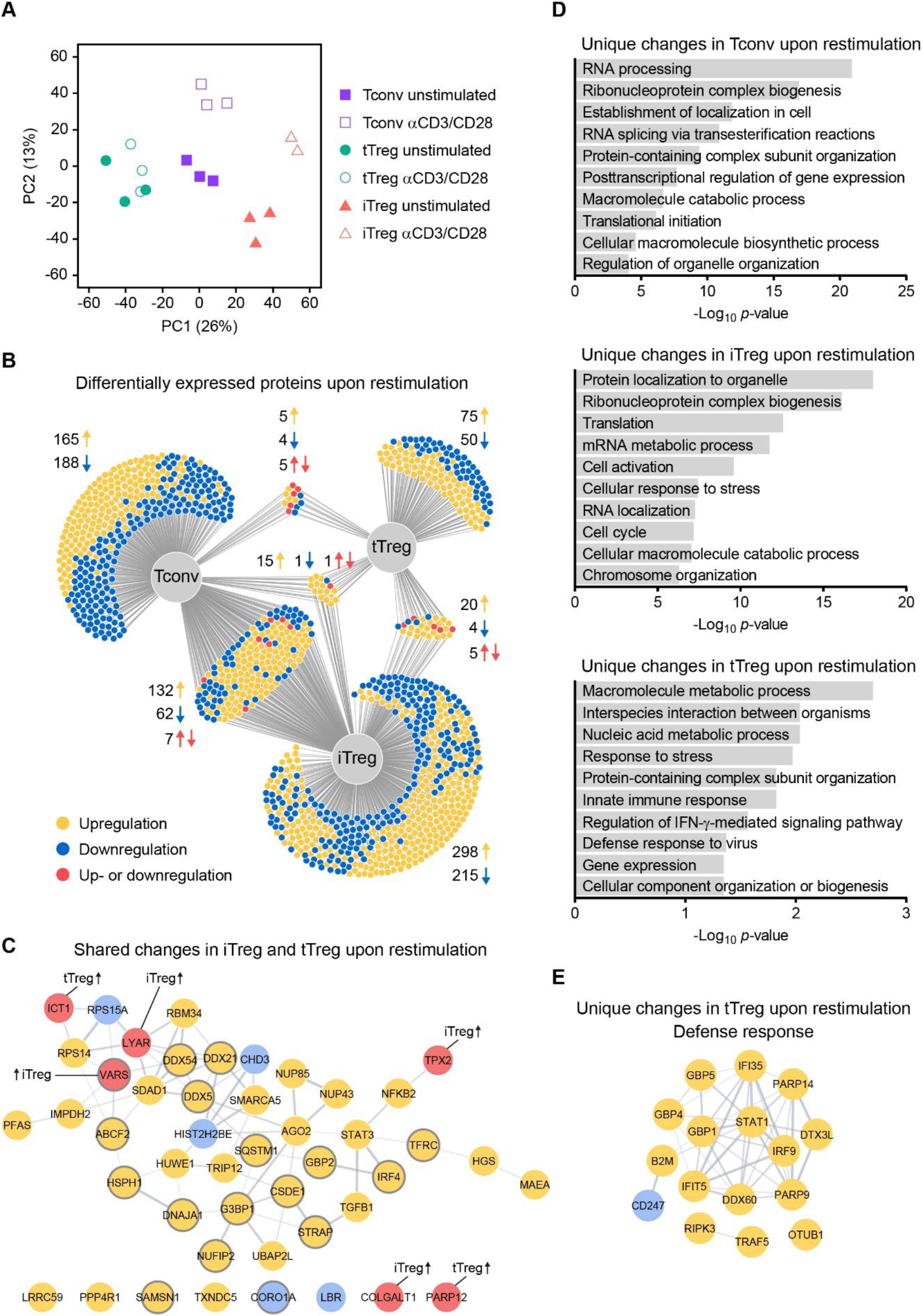
Proteomic analysis of CD3/CD28-stimulated Tconv, tTreg and iTreg cells. **(A)** PCA plot of the proteome of Tconv, tTreg and iTreg cells that were restimulated or not (*n* = 3, *n* = 2 only for CD3/CD28-stimulated iTreg cells). **(B)** DiVenn diagram showing shared and unique proteins that were differentially expressed upon CD3/CD28 stimulation in Tconv, tTreg and iTreg cells (two-sided *t*-test, *p* < 0.05). Upregulated proteins (yellow), downregulated proteins (blue) or proteins of which the expression changed in opposite directions (red) are shown. **(C)** STRING network of proteins that changed expression in both iTreg and tTreg cells upon restimulation. Proteins that also changed expression in Tconv cells are displayed as nodes with a grey border. Upregulated or downregulated proteins are depicted as yellow or blue nodes, respectively. Red nodes indicate proteins of which the expression changed in opposite directions, annotated with the cell type that showed upregulation. **(D)** GO enrichment analyses of unique differentially expressed proteins in restimulated Tconv, tTreg or iTreg cells, showing ten significantly enriched GO biological processes per cell type. **(E)** STRING analysis of proteins that uniquely changed expression in restimulated tTreg cells, showing a network of proteins involved in an interferon-mediated defense response.

Linear regression analysis revealed that the response of Tconv and iTreg cells correlated more than the response of tTreg and iTreg cells (**Supplementary Figure 4B**). Accordingly, a DiVenn diagram revealed over 200 proteins that commonly altered their expression in both Tconv and iTreg cells (**Figure 4B**). These proteins are primarily involved in cell activation, macromolecular biosynthesis and responses to stress according to GO analysis (**Supplementary Figure 4C**). 46 proteins commonly altered their expression in tTreg and iTreg cells, of which 17 also altered their expression in Tconv cells (**Figure 4B**). These proteins are involved in a variety of cellular processes, including lymphocyte activation (**Figure 4C**). For many proteins, however, levels were uniquely altered in either Tconv, iTreg or tTreg cells. These proteins are involved in metabolism, RNA processing and biogenesis (**Figure 4D**). Noteworthy, tTreg cells uniquely upregulated proteins involved in an interferon-mediated defense response (**Figure 4E**). In conclusion, tTreg cells showed a relatively modest response to CD3/CD28-mediated activation as compared to iTreg and Tconv cells. Furthermore, all three cell types responded largely in a unique fashion, in particular regarding processes related to macromolecular biosynthesis.

### 3.5 A core signature reflecting unique Treg cell properties is present in tTreg but not iTreg cells

Recently, we established that *ex vivo* naïve and effector Treg cells from human peripheral blood have a common proteomic signature that distinguishes them from CD4^+^ Tconv cells, which we termed the Treg cell core proteomic signature (22). This signature contains proteins with Treg cell-specific high or low expression reflecting unique properties regarding signal transduction, transcriptional regulation, iron storage, lysosomal processes and cell metabolism (**Figure 5A**, **B**). Here, we questioned whether the Treg cell core signature is also present in iTreg cells. In our proteomics dataset, we detected 46 out of 51 proteins of the Treg cell core signature and evaluated their expression in expanded Tconv, tTreg and iTreg cells. PCA indicated distinct clustering of these three cell types based on Treg cell core signature protein levels, with iTreg cells clearly clustering away from tTreg cells (**Figure 5C**). Hierarchical clustering based on this signature also separated the three cell types (**Figure 5D**). The Treg cell core signature, with some exceptions, was preserved in tTreg cells following *in vitro* expansion, as reported previously (22) (**Figure 5D**). tTreg and iTreg cells shared a number of the Treg cell core signature proteins with characteristic high (cluster 1—including FOXP3, as expected) or low (cluster 3) expression levels relative to Tconv cells. Moreover, iTreg cells, as opposed to *in vitro*- expanded tTreg cells, expressed the core signature proteins MVP, HK1, ME2 and APOO at characteristic levels relative to those in Tconv cells. Nevertheless, 26 out of 46 Treg cell core signature proteins did not exhibit the characteristic expression pattern (compared to Tconv cells) in iTreg cells. For instance, iTreg cells expressed high levels of the transcription factor STAT4, while low levels in Treg cells are important to conserve Treg cell function (22). These proteomic data suggest that iTreg cells share only a part of the unique characteristics of *ex vivo* Treg cells.

**Figure 5.**
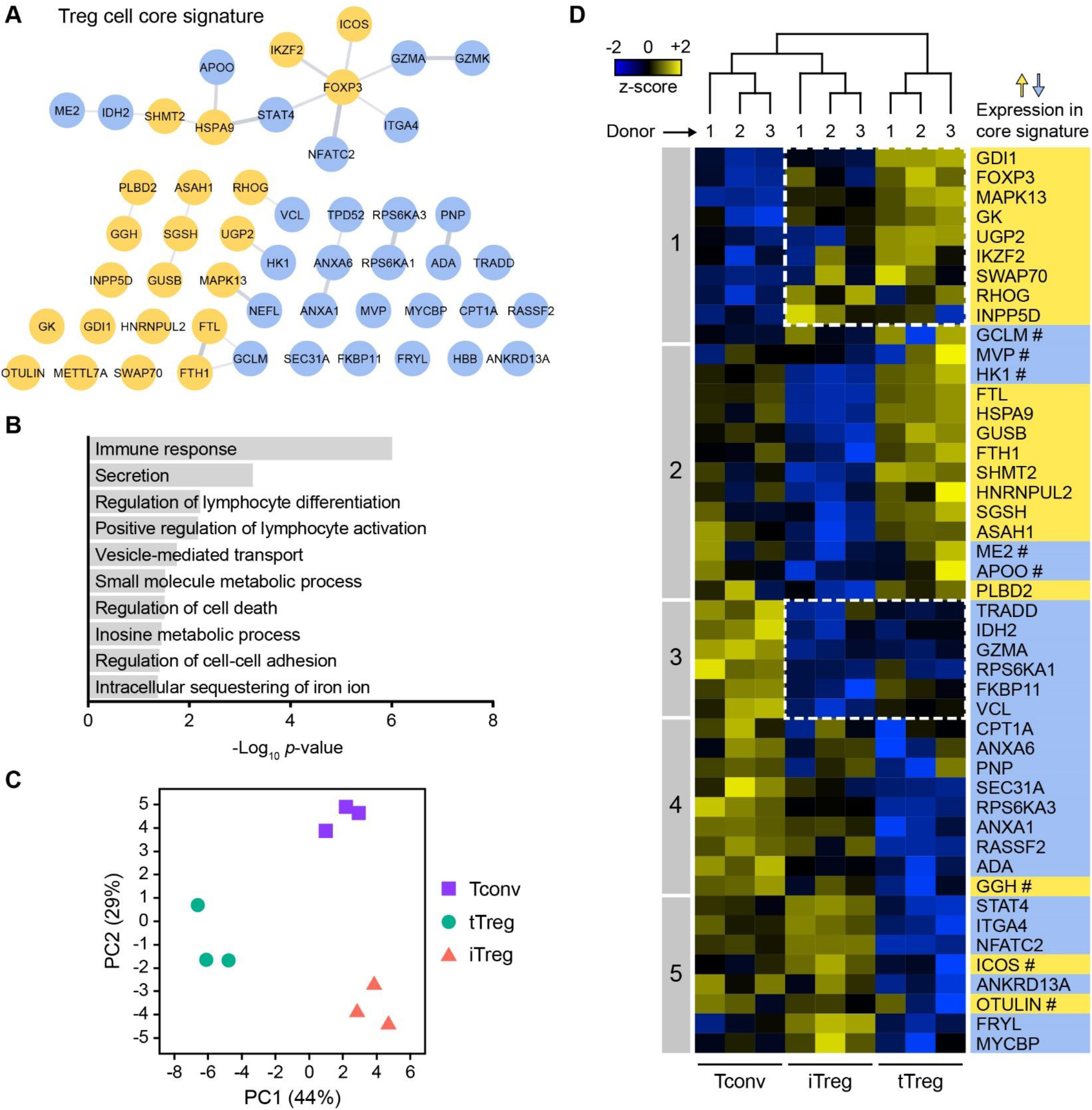
A Treg cell core signature is present in *in vitro*-expanded tTreg but not iTreg cells. **(A)** STRING network displaying 51 proteins of a previously defined Treg cell core proteomic signature (22). Upregulated and downregulated proteins are shown as yellow and blue nodes, respectively. **(B)** GO enrichment analysis of the Treg cell core signature proteins, showing ten significantly enriched GO biological processes. **(C)** PCA plot based on the expression of Treg cell core signature proteins in unstimulated Tconv, tTreg and iTreg cells (*n* = 3). **(D)** Heat map showing hierarchical clustering of 46 detected proteins of the Treg cell core signature in unstimulated Tconv, tTreg and iTreg cells. Z-scores depicting relative protein expression values are color-coded. Numbered boxes (1–5) indicate clusters of proteins. Protein names are colored according to high (yellow) or low (blue) relative expression in the Treg cell core signature. Proteins are marked (#) when the expression in the expanded and rested tTreg cells used here deviates from the described core signature (22).

### 3.6 iTreg cells do not acquire the Treg cell core signature upon CD3/CD28-mediated activation

Since iTreg cells responded greatly to CD3/CD28-mediated restimulation in terms of altered protein expression, we determined whether they obtained more of the Treg cell core signature under these conditions. The core signature is present in both naïve and effector tTreg cells (22), indicating that core signature proteins in tTreg cells do not respond majorly to cell activation. This was indeed observed in a PCA, where levels of Treg cell core signature proteins did not alter in tTreg cells upon CD3/CD28-mediated restimulation (**Figure 6A**). Interestingly, the levels of these proteins also did not alter much in Tconv or iTreg cells upon restimulation (**Figure 6A**). Accordingly, hierarchical clustering showed grouping of core protein expression levels based on cell type, with tTreg cells being most distinct from Tconv and iTreg cells (**Figure 6B**). Part of the Treg cell core signature was shared by iTreg and tTreg cells, including proteins with either high (cluster 2) or low (cluster 3) expression levels compared to Tconv cells. Furthermore, core signature proteins HK1, APOO, ME2 and FKBP11 were expressed at characteristic low levels in iTreg cells (cluster 1). However, the major part of the Treg cell core signature proteins did not show characteristic expression levels in iTreg cells. Differential expression of core signature proteins NFATC2, ITGA4 and ICOS between tTreg and iTreg cells was confirmed by flow cytometry (**Figure 6C**, **Supplementary Figure 5**). Low expression of NFATC2 is a critical feature of Treg cell function (22), whereas iTreg cells shared high expression with Tconv cells. In conclusion, Treg cell core signature protein expression in iTreg cells was not more pronounced after CD3/CD28-mediated activation.

**Figure 6.**
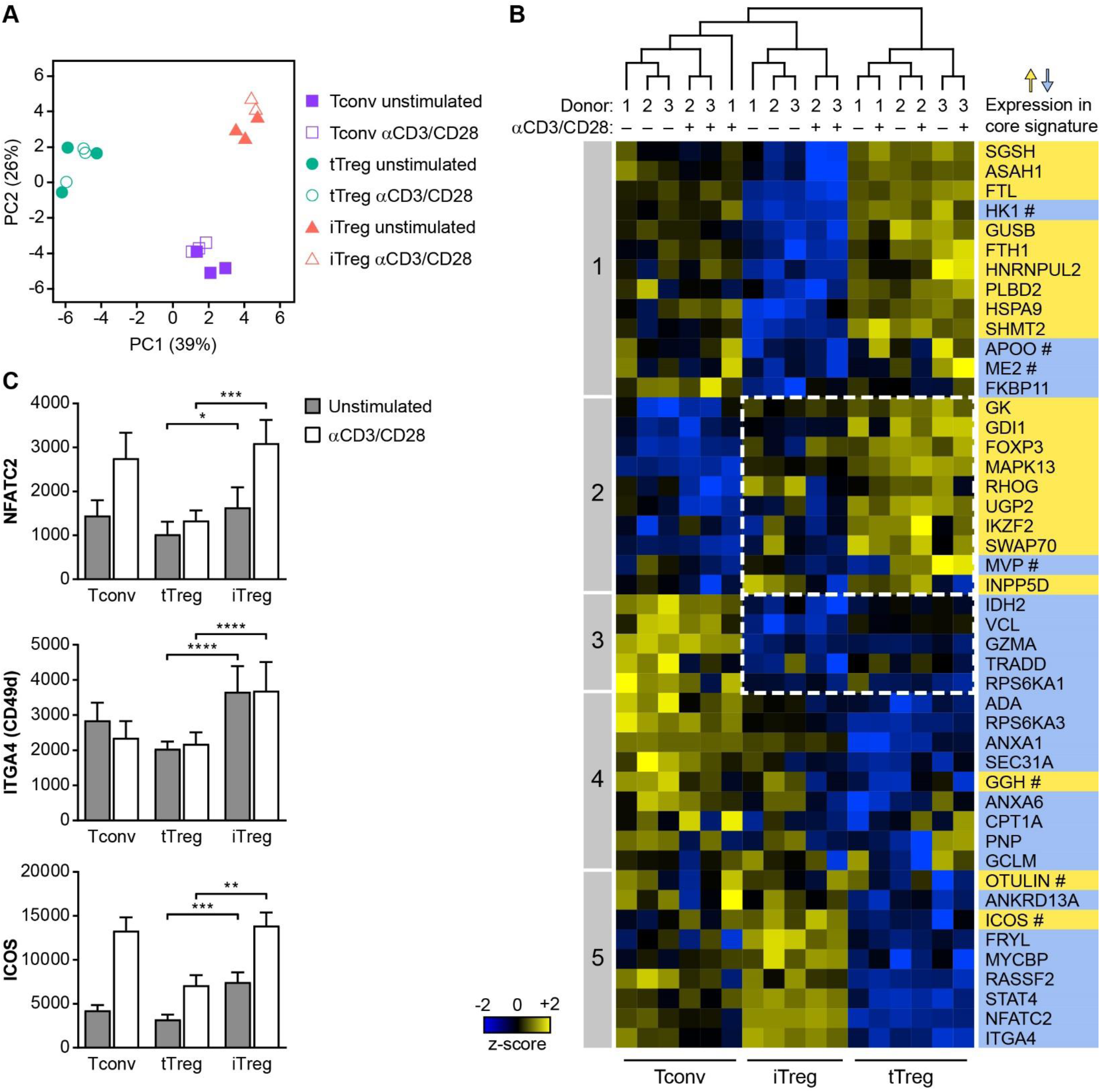
Expression levels of Treg cell core signature proteins do not change in any cell type upon CD3/CD28-mediated restimulation. **(A)** PCA plot based on the expression of Treg cell core signature proteins in Tconv, tTreg and iTreg cells that were restimulated or not (*n* = 3, *n* = 2 only for CD3/CD28-stimulated iTreg cells). **(B)** Heat map showing hierarchical clustering of 46 detected proteins of the Treg cell core signature in Tconv, tTreg and iTreg cells that were CD3/CD28-stimulated or not. Z-scores depicting relative protein expression values are color-coded. Numbered boxes (1–5) indicate clusters of proteins. Protein names are colored according to high (yellow) or low (blue) relative expression in the Treg cell core signature. Marking (#) indicates the same as in Figure 5D. **(C)** Quantification of flow cytometric analysis of the protein expression of total NFATC2 (*n* = 4), surface ITGA4 (CD49d) (*n* = 5) and surface ICOS (*n* = 5) on indicated cell populations. MFI is depicted on the y-axis. Two-way ANOVA with Tukey’s *post hoc* test was used for statistical analysis. Data are presented as mean ± SEM. Sample size (*n*) represents cells from individual donors, analyzed in independent experiments. \**p*< 0.05, \*\**p* < 0.01, \*\*\**p* < 0.001, \*\*\*\**p* < 0.0001.

### 3.7 iTreg cells acquire aspects of an effector Treg cell signature

Besides the Treg cell core signature, we previously observed a unique set of proteins that was characteristically expressed at relatively high or low levels in *ex vivo* effector Treg cells, which sets them apart from Tconv cells and naïve Treg cells (22). This was termed the effector Treg cell signature, of which the highly expressed proteins are predominantly associated with DNA replication (e.g. MCM proteins), apoptosis (e.g. FAS, CASP3), vesicular trafficking (e.g. STXBP2, NSF) and cytoskeletal reorganization (e.g. TUBA1B, TUBB) (**Figure 7A**). Lowly expressed proteins of the effector Treg cell signature primarily included those associated with signaling by the TCR/CD3 complex and CD28 (e.g. THEMIS, TRAT1, VAV1), phosphatidylinositol (e.g. INPP4A, INPP4B), NF-κB (e.g. NFKB1, NFKB2) and JAK-STAT (e.g. STAT3, IL7R) (**Figure 7B**). Here, we examined the effector Treg cell signature in unstimulated and restimulated Tconv, tTreg and iTreg cells and questioned whether iTreg cells acquire effector Treg cell-like properties.

**Figure 7.**
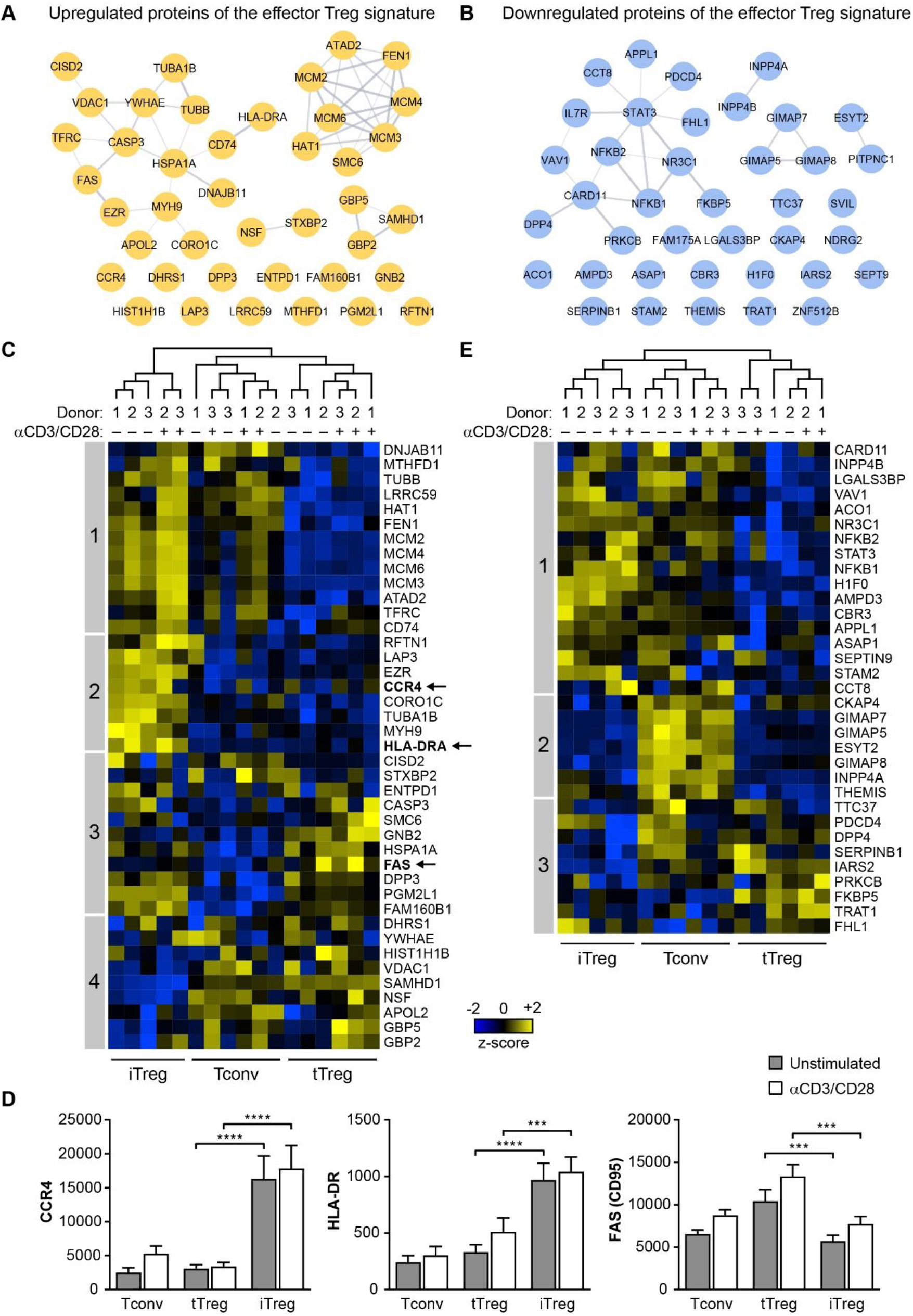
Aspects of an effector Treg cell signature are present in iTreg cells. **(A, B)** STRING networks showing 41 upregulated **(A)** or 39 downregulated proteins **(B)** of a previously defined effector Treg cell proteomic signature (22) as yellow and blue nodes, respectively. **(C)** Heat map showing hierarchical clustering of 41 detected proteins of the upregulated part of the effector Treg cell signature in Tconv, tTreg and iTreg cells that were restimulated or not (*n* = 3, *n* = 2 for CD3/CD28-stimulated iTreg cells). Z-scores showing relative protein expression values are color-coded. Numbered boxes (1–4) indicate clusters of proteins. CCR4, HLA-DRA and FAS are marked, as these molecules are also analyzed in **D**. **(D)** Quantification of flow cytometric analysis of the protein expression of surface CCR4 (*n* = 4), surface HLA-DR (*n* = 4) and surface FAS (CD95) (*n* = 4) on indicated cell populations. MFI is depicted on the y-axis. Two-way ANOVA with Tukey’s *post hoc* test was used for statistical analysis. Data are presented as mean ± SEM. Sample size (*n*) represents cells from individual donors, analyzed in independent experiments. \*\*\**p* < 0.001, \*\*\*\**p* < 0.0001. **(E)** Heat map showing hierarchical clustering of 33 detected proteins of the downregulated part of the effector Treg cell signature in Tconv, tTreg and iTreg cells that were restimulated or not (*n* = 3, *n* = 2 for CD3/CD28-stimulated iTreg cells). Z-scores showing relative protein expression values are color-coded. Numbered boxes (1–3) indicate clusters of proteins.

We detected all 41 signature proteins with high expression levels in effector Treg cells (effector Treg^hi^). Hierarchical clustering based on expression of these 41 signature proteins showed that iTreg cells clustered away from Tconv and tTreg cells (**Figure 7C**). iTreg cells, as opposed to tTreg cells, either before or after CD3/CD28-mediated activation, showed enrichment of the effector Treg^hi^ signature with 21 out of 41 proteins exhibiting relatively high expression (clusters 1–2) (**Figure 7C**, **Supplementary Table 3**). These proteins were primarily involved in DNA replication (e.g. MCM proteins) and cytoskeletal processes such as cytokinesis (e.g. MYH9, CORO1C), indicating cell proliferation. Additionally, the majority of 11 proteins in cluster 3 indicated relatively high expression of effector Treg^hi^ signature proteins in both iTreg and tTreg cells but not Tconv cells. Proteins in this cluster included CD39 (ENTPD1), a well-known effector molecule on Treg cells, which catalyzes a critical step in the conversion of ATP into immunosuppressive adenosine (44). Flow cytometry verified that CCR4 and HLA-DR had highest expression on iTreg cells, that CD39 was expressed on both iTreg and tTreg cells (**Figure 2**), whereas FAS was highly expressed on tTreg cells, all in agreement with the results from the proteome measurements (**Figure 7D**, **Supplementary Figure 6**). Thus, altogether, 32 out of 41 Treg^hi^ signature proteins were also expressed at relatively high levels in iTreg cells.

We detected 33 out of 39 signature proteins with characteristic low relative expression levels in effector Treg cells (effector Treg^lo^) (**Figure 7E**, **Supplementary Table 3**). Hierarchical clustering based on expression of these proteins showed that iTreg cells clustered away from Tconv and tTreg cells (**Figure 7E**). Among the 33 effector Treg^lo^ signature proteins, 16 exhibited relatively low expression in iTreg cells (clusters 2–3). Proteins in cluster 2 were also expressed at low levels in tTreg cells and included INPP4A, THEMIS and members of the GIMAP family. Another group of proteins was mainly expressed at low levels in iTreg cells, but not in tTreg cells (cluster 3). However, 17 out of 33 effector Treg^lo^ signature proteins did not have the characteristic low expression in iTreg cells, whereas they were lowly expressed in tTreg cells (cluster 1). These included proteins that play a role in signaling downstream of immune receptors, such as NFKB1, NFKB2, CARD11, STAT3, VAV1, STAM2 and INPP4B. These data show that the effector Treg cell signature is partially present in iTreg cells, indicating that iTreg cells resemble *ex vivo* effector Treg cells to some extent.

## 4 Discussion

The Treg cell lineage can be subdivided into tTreg and pTreg cells based on their developmental origin (15, 47). tTreg cells have escaped negative selection in the thymus and are mainly geared towards self-antigen recognition (15). They comprise the majority of Treg cells *in vivo* and control systemic and tissue-specific autoimmunity (48, 49). pTreg cells recognize foreign antigen and are largely restricted to mucosal surfaces and the maternal-fetal interface (14, 50). Both cell types can be distinguished in mice, where common and unique features in their phenotypes and transcriptome profiles have been observed (11, 12). tTreg and pTreg cells complement each other in immune homeostasis (51, 52), which may in part be due to their distinct TCR repertoires and differential expression of certain transcriptional regulators (14, 53). In human, for lack of surface markers, effector phenotype tTreg or pTreg cells cannot be discriminated, which complicates the interpretation of many human Treg cell studies. Moreover, iTreg cells are often employed as an *in vitro* model for Treg cell function, differentiation or metabolism, which increases the complexity of Treg cell literature.

Here, we questioned to what extent iTreg cells resemble tTreg cells in their protein expression, focusing on conserved features of *ex vivo* Treg cells as previously defined (22). We performed a side-by-side analysis of *in vitro*-generated iTreg cells with tTreg cells isolated to high purity from the blood and also included Tconv cells as a reference. Proteomics gives more direct insight into the functional cellular programs than transcriptomics, since protein and transcript levels often differ due to regulation at protein and mRNA levels (22, 54). Our proteomic analysis pointed out that iTreg cells display a protein expression profile that markedly sets them apart from tTreg cells.

Among the proteins with a relatively low expression in iTreg cells compared to tTreg cells in the steady state, we found an enrichment of proteins involved in a variety of metabolic processes. iTreg cells are frequently used to study Treg cell metabolism and reportedly utilize different metabolic programs than other CD4^+^ T cell subsets (26, 27, 55). Our findings now indicate dissimilarity between iTreg and tTreg cells in terms of metabolism, suggesting that iTreg cells do not mirror (t)Treg cells found *in vivo*. However, it remains to be resolved to what extent iTreg cells share metabolic features with pTreg cells, which also derive from Tconv cells under the influence of TGF-β (14).

We also observed a substantial number of proteins with relatively high expression in iTreg cells compared to tTreg cells in the steady state. Notable was the enrichment of signal transduction proteins involved in lymphocyte activation. This different configuration of signaling molecules likely influences the potential response of iTreg and tTreg cells to extracellular stimuli. Indeed, CD3/CD28-mediated activation of these cells resulted in unique responses of different magnitudes. Whereas iTreg cells strongly responded by differential expression of a large number of proteins, tTreg cells showed only a modest response. Adaptations in signaling pathways downstream of immune receptors such as the TCR and CD28 may underlie the attenuated signal transduction capacity in tTreg cells, which has been shown for *ex vivo* Treg cells before (22). iTreg cells shared more features with Tconv cells than with tTreg cells, which are associated with metabolism, responses to stress and lymphocyte activation. Like iTreg cells, Tconv cells responded strongly to CD3/CD28-mediated activation, yet both cell types responded largely in a unique manner.

The proteomic similarities between iTreg and tTreg cells were rather limited. GO analysis only indicated an enrichment of proteins involved in endoplasmic reticulum-associated processes. We confirmed that iTreg cells acquire Treg cell-like features such as increased expression of FOXP3 and CTLA-4 (24), especially following CD3/CD28-mediated activation. FOXP3 expression in iTreg cells was heterogeneous, as has been observed by others (26, 27, 55). We also observed shared high or low expression of IKZF3 and YY1, respectively, which can interact with FOXP3 to regulate transcription (45, 46). iTreg cells did not express Helios at similar high levels found in tTreg cells, but minor upregulation was observed. It has been demonstrated in mice that, after recent egress from the thymus, a small proportion of naïve CD4^+^ Tconv cells has the capacity to develop into Helios^+^ pTreg cells (56). Nevertheless, Helios is preferentially expressed in tTreg cells and marks stable commitment to the human Treg cell lineage (7, 8).

Stable expression of FOXP3 is established at the epigenetic level by demethylation of several regions within the *FOXP3* gene locus. A critical region for sustained FOXP3 expression is the TSDR in the CNS2 enhancer, which we confirmed to be highly demethylated in tTreg cells but not in iTreg cells, as described (5, 15). However, a recent study comprehensively investigated mouse iTreg cells at the epigenetic level after generation in the presence or absence of CD28 costimulation (57), as strong CD28 signaling reportedly hampers iTreg cell development (58). iTreg cells deprived of CD28 costimulation showed increased DNA demethylation within Treg cell-specific genes, leading to more stable Foxp3 expression and enhanced suppressive function, which even persisted after adoptive transfer in mice (57).

We asked whether iTreg cells resemble Treg cells that are found *in vivo*. tTreg cells relocate from lymphoid organs to peripheral tissues, where they are under the continuous influence of environmental signals. They have a degree of plasticity and can adapt themselves to locally resident effector T cells by responding to specific cytokines (49). This involves the gain of lineage-determining transcription factors such as GATA-3 or T-bet in addition to FOXP3, which alters their functionality. However, tTreg cells retain core features that safeguard their suppressive nature and prevent them from exerting pro-inflammatory functions (22). To establish whether iTreg cells acquire such features, we used proteomic signatures from human Treg cells that were analyzed directly from peripheral blood (22).

The first benchmark we used was the Treg cell core signature, a conserved set of proteins between naïve and effector Treg cells compared to other CD4^+^ T cell subsets that emphasizes critical Treg cell aspects (22). This signature strongly suggests differential wiring of signaling pathways, given the constitutive differential expression of key signal transduction components between Treg and Tconv cells. We observed that iTreg cells only share a limited part of these conserved characteristics, while the signature was largely present in tTreg cells, regardless of whether the cells were resting or had been restimulated via CD3/CD28. As iTreg cells do not possess the signal transduction profile of the Treg cell core signature, this likely leads to differential wiring and a different response to stimuli.

The Treg cell core signature includes low expression of STAT4 and NFATC2, which are mediators of inflammatory cytokine expression (59, 60). Low expression of these molecules may aid Treg cells to protect their anti-inflammatory nature in a pro-inflammatory environment, while they can still respond to pro-inflammatory cytokines through other pathways (22). It has been demonstrated that deliberate elevation of STAT4 expression in tTreg cells increases their susceptibility to pro-inflammatory cytokines, provoking effector cytokine production and loss of FOXP3 expression (22). Both STAT4 and NFATC2 had relatively high expression in iTreg cells compared to tTreg cells, suggesting that iTreg cells are more susceptible to respond to pro-inflammatory cues in a pro-inflammatory manner, such as having the capacity to produce IFN-γ (61). In addition to the fragile epigenetic stability of iTreg cells, our findings now suggest that the susceptibility of iTreg cells to abandoning the Treg cell fate upon exposure to pro-inflammatory cytokines may also be caused by their high expression of involved mediators, such as STAT4 and NFATC2.

The second benchmark we examined was an effector Treg cell signature, which sets effector Treg cells apart from naïve Treg cells and other CD4^+^ T cell subsets (22). Relative to protein expression in tTreg and Tconv cells, iTreg cells acquired aspects of the effector Treg cell signature, indicating that iTreg cells may resemble *ex vivo* effector Treg cells to some extent. The enrichment of the effector Treg cell signature in iTreg cells may fit with the idea that the effector Treg cell population contains both activated tTreg cells and pTreg cells (62, 63).

We propose that the core and effector Treg cell proteomic signatures (22) are useful reference tools to define Treg cell populations. In conclusion, our global proteomic analysis showed that iTreg and tTreg cells each share more protein expression with Tconv cells than with one another. Moreover, part of the overall protein expression profile of iTreg cells is unique. Thus, despite the acquisition of certain Treg cell-like features, the conversion of Tconv cells into iTreg cells generates cells that become even more distinct from tTreg cells than Tconv cells are. Therefore, we propose that experiments in which iTreg cells—cultured in commonly used conditions as in the present article—are employed as a model for studying Treg cell differentiation, function or metabolism should be interpreted with caution.

## 5 Data availability

Proteomics data on tTreg and Tconv cells have been deposited as outlined before (22). Proteomics data on iTreg cells will be deposited before publication of this manuscript.

## 6 Conflict of interest

The authors declare that the research was conducted in the absence of any commercial or financial relationships that could be construed as a potential conflict of interest.

## 7 Author contributions

MM, SdK and JB designed the study and wrote the manuscript, MM and SdK performed experiments and analyzed the data, ES provided technical assistance, EC provided data and insights, MvdB performed mass spectrometry measurements, DA provided conceptual advice in the study design and contributed to the manuscript.

## 8 Funding

This study was supported by Oncode, ZonMW-TOP grant 40-00812-98-13071, and grants ICI-00014 and ICI-00025 from the Institute for Chemical Immunology, funded by ZonMW Gravitation, and Strategic Funds from the LUMC.

## Acknowledgements

We thank the staff members of the flow cytometry facility at the Netherlands Cancer Institute and the flow cytometry core facility at Leiden University Medical Center for their technical assistance.

---

**Supplementary Figure 1.**
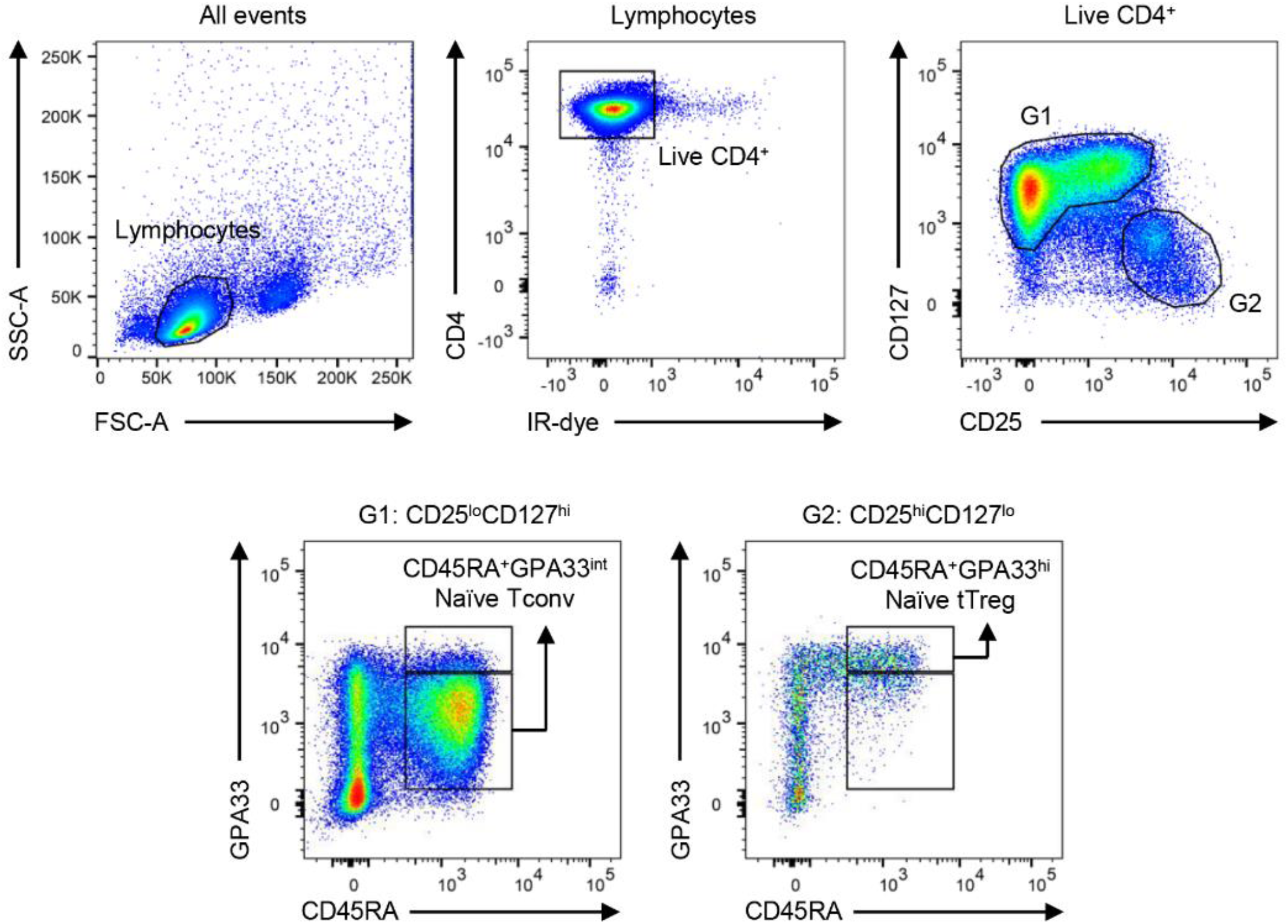
Gating strategy to sort naïve Tconv and naïve tTreg cells from human peripheral blood by flow cytometry. Pre-enriched total CD4^+^ T cells from human peripheral blood were first separated into live CD4^+^CD25^lo^CD127^hi^ cells within gate 1 (G1) and live CD4^+^CD25^hi^CD127^lo^ cells within gate 2 (G2). Subsequently, CD45RA^+^GPA33^int^ naïve Tconv cells and CD45RA^+^GPA33^hi^ naïve tTreg cells were selected within G1 and G2, respectively.

**Supplementary Figure 2.**
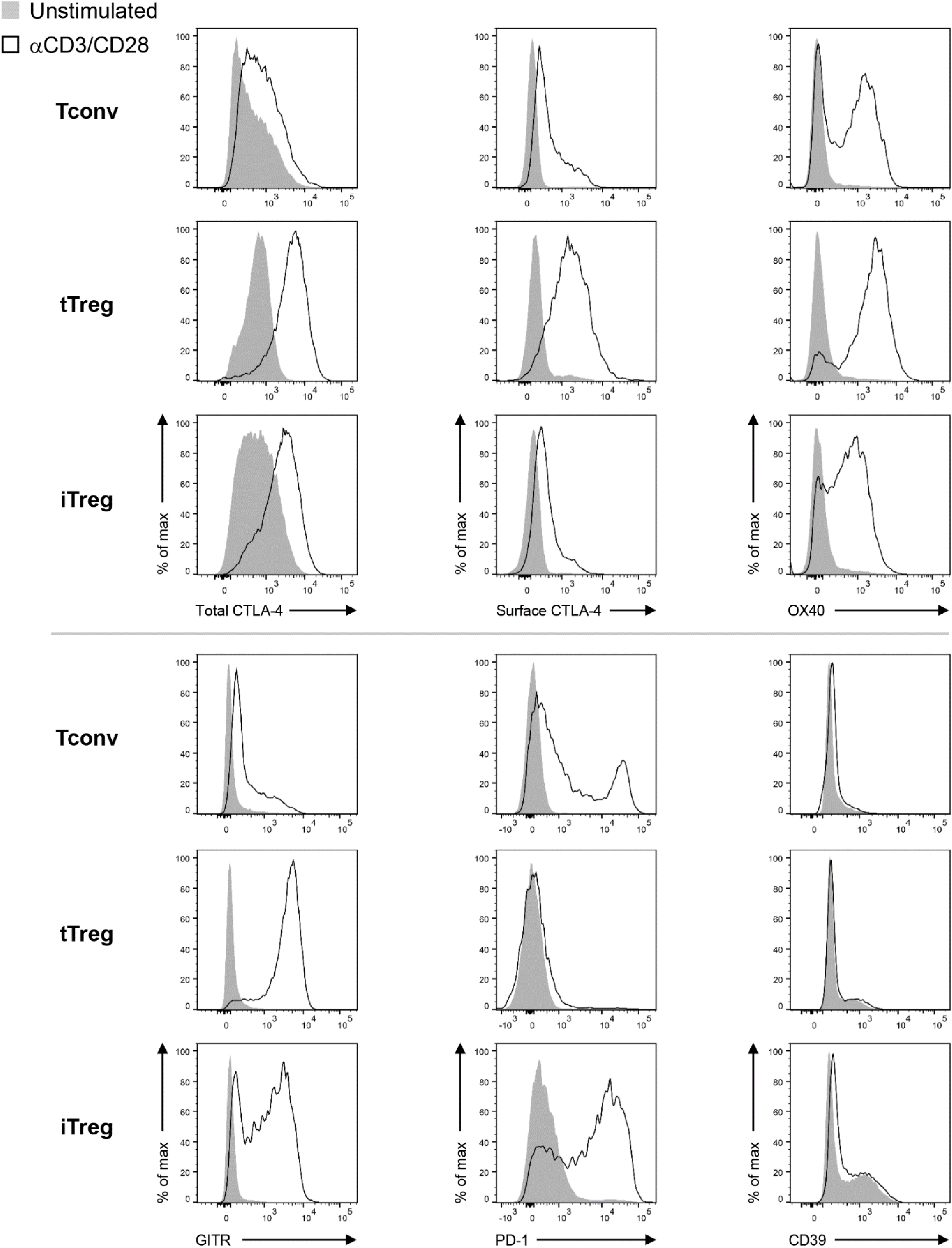
Representative flow cytometry histograms showing protein expression of total CTLA-4 (*n* = 13), surface CTLA-4 (*n* = 8), surface OX40 (*n* = 8), surface GITR (*n* = 8), surface PD-1 (*n* = 8) and surface CD39 (*n* = 8) on Tconv, tTreg and iTreg cells that were restimulated via CD3/CD28 or not.

**Supplementary Figure 3.**
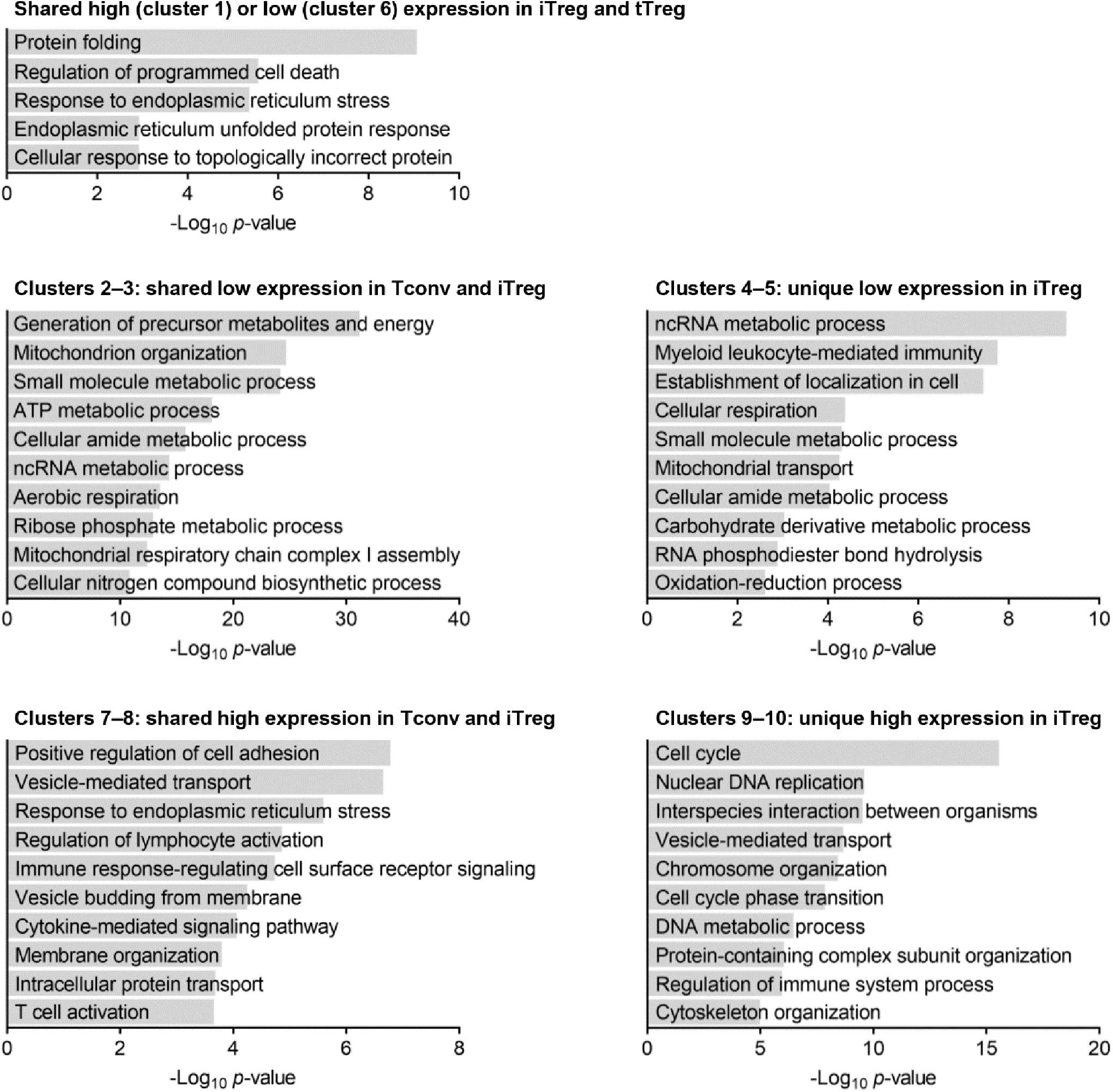
GO biological processes enrichment analyses of indicated clusters of proteins from Figure 3B that were differentially expressed between unstimulated Tconv, tTreg and iTreg cells (*p* < 0.05).

**Supplementary Figure 4.**
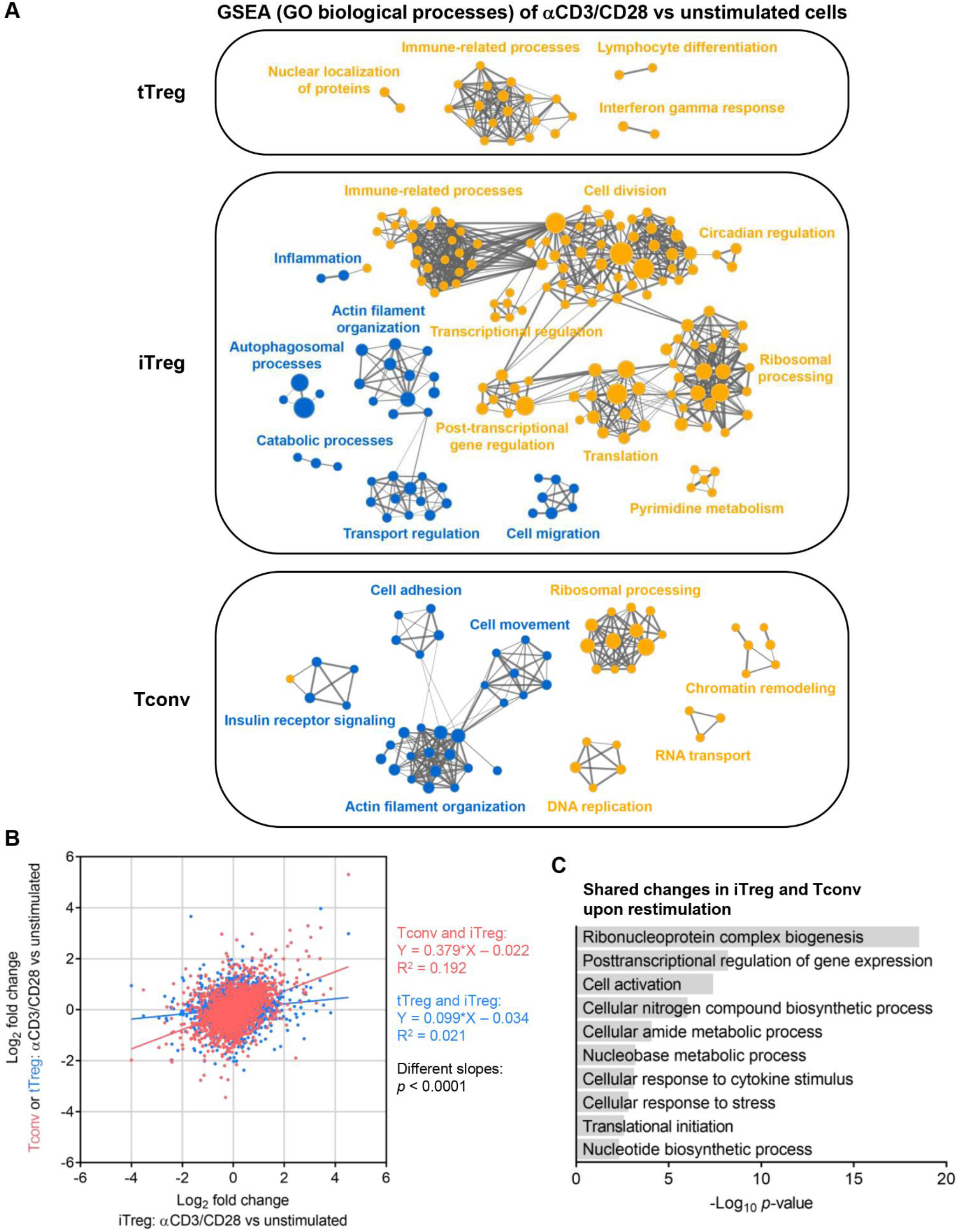
Analysis of the proteomic responses of tTreg, iTreg and Tconv cells to CD3/CD28-mediated restimulation. **(A)** GSEA enrichment maps of the proteomic responses of tTreg, iTreg or Tconv cells to CD3/CD28 restimulation, showing enriched GO biological processes in yellow (upregulated) and blue (downregulated). Gene sets are depicted as nodes, of which the size represents the number of genes in the gene set. Gene sets sharing a large number of genes are clustered and are connected by lines of which the thickness corresponds with the number of shared genes. Clusters of similar biological processes are annotated using general terms. **(B)** Linear regression analysis of the proteomic response of iTreg cells (x-axis) compared to the responses of either Tconv (red, y-axis) or tTreg (blue, y-axis) cells, revealing different regression lines (*p* < 0.0001). **(C)** GO biological processes enrichment analysis of shared changes in iTreg and Tconv cells, but not tTreg cells, upon CD3/CD28-mediated activation (*p* < 0.05).

**Supplementary Figure 5.**
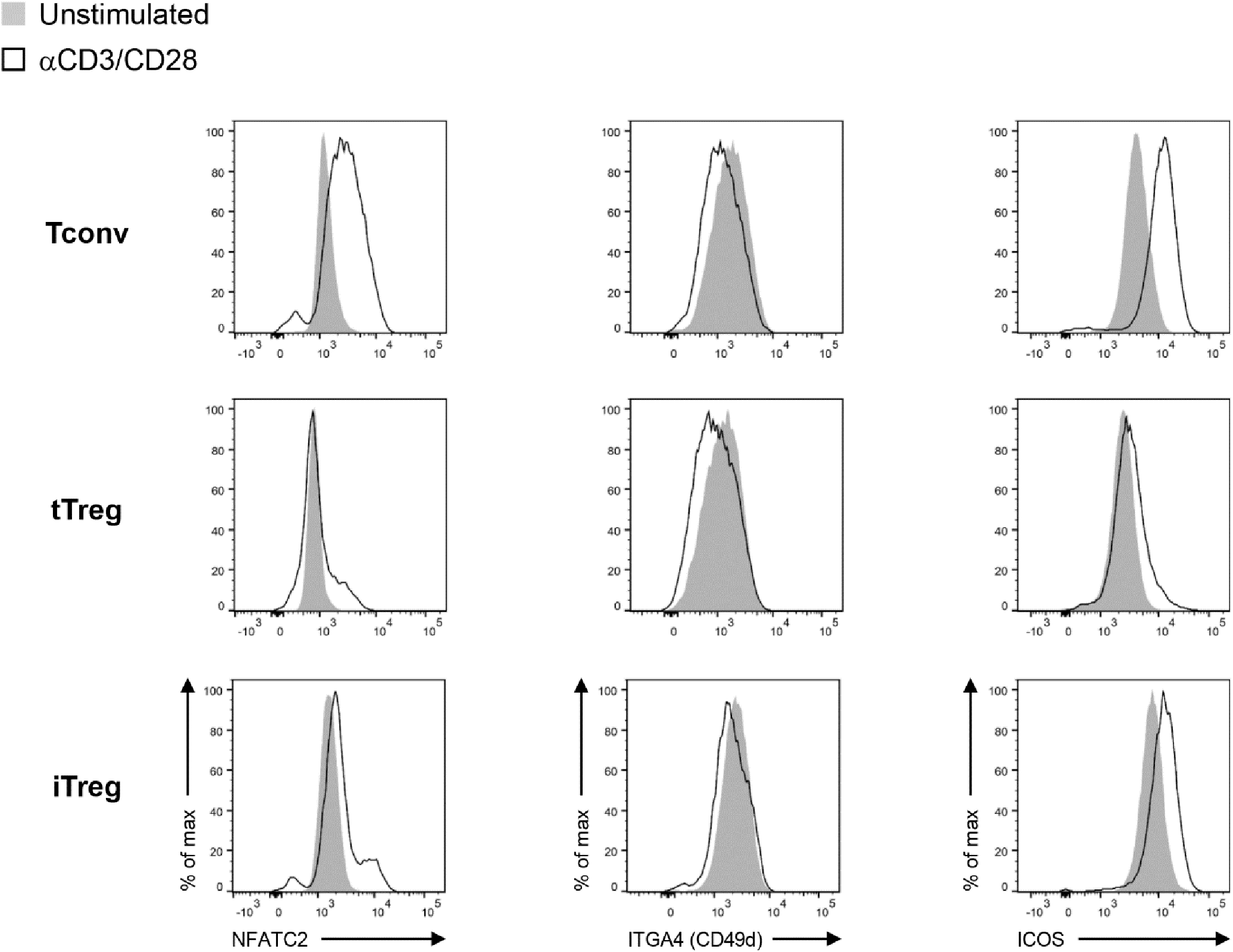
Representative flow cytometry histograms showing protein expression of total NFATC2 (*n* = 4), surface ITGA4 (CD49d) (*n* = 5) and surface ICOS (*n* = 5) on Tconv, tTreg and iTreg cells that were restimulated via CD3/CD28 or not.

**Supplementary Figure 6.**
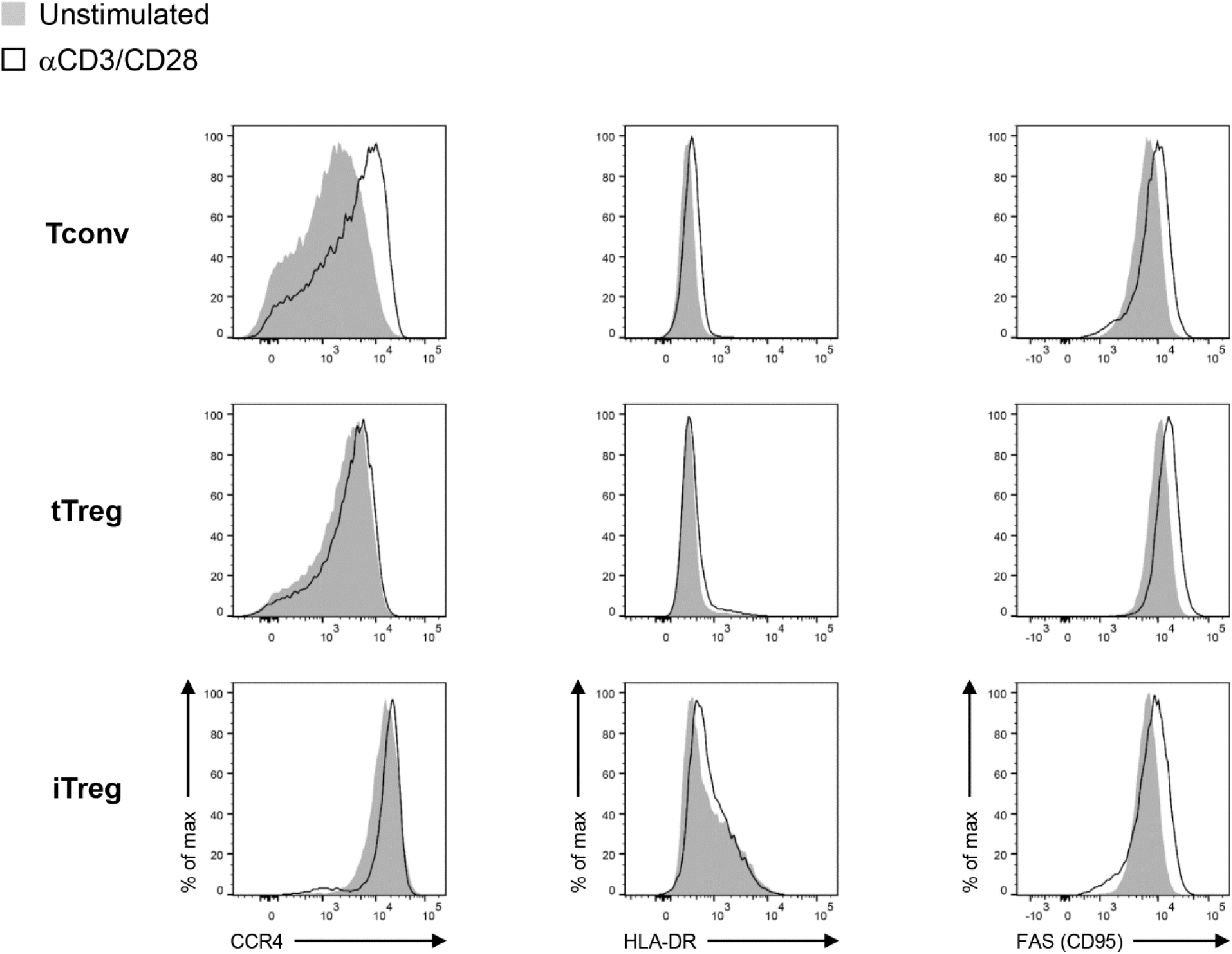
Representative flow cytometry histograms showing protein expression of surface CCR4 (*n* = 4), surface HLA-DR (*n* = 4) and surface FAS (CD95) (*n* = 4) on Tconv, tTreg and iTreg cells that were restimulated via CD3/CD28 or not.

## Notes

### Competing Interest Statement

The authors have declared no competing interest.

